# A Deep Learning Framework for Predicting Human Essential Genes by Integrating Sequence and Functional data

**DOI:** 10.1101/2020.08.04.236646

**Authors:** Xue Zhang, Weijia Xiao, Wangxin Xiao

## Abstract

Essential genes are necessary to the survival or reproduction of a living organism. The prediction and analysis of gene essentiality can advance our understanding to basic life and human diseases, and further boost the development of new drugs. Wet lab methods for identifying essential genes are often costly, time consuming, and laborious. As a complement, computational methods have been proposed to predict essential genes by integrating multiple biological data sources. Most of these methods are evaluated on model organisms. However, prediction methods for human essential genes are still limited and the relationship between human gene essentiality and different biological information still needs to be explored. In addition, exploring suitable deep learning techniques to overcome the limitations of traditional machine learning methods and improve the prediction accuracy is also important and interesting. We propose a deep learning based method, DeepSF, to predict human essential genes. DeepSF integrates sequence features derived from DNA and protein sequence data with features extracted or learned from different types of functional data, such as gene ontology, protein complex, protein domain, and protein-protein interaction network. More than 200 features from these biological data are extracted/learned which are integrated together to train a cost-sensitive deep neural network by utilizing multiple deep leaning techniques. The experimental results of 10-fold cross validation show that DeepSF can accurately predict human gene essentiality with an average AUC of 95.17%, the area under precision-recall curve (auPRC) of 92.21%, the accuracy of 91.59%, and the F1 measure about 78.71%. In addition, the comparison experimental results show that DeepSF significantly outperforms several popular traditional machine learning models (SVM, Random Forest, and Adaboost), and performs slightly better than a recent deep learning model (DeepHE). We have demonstrated that the proposed method, DeepSF, is effective for predicting human essential genes. Deep learning techniques are promising at both feature learning and classification levels for the task of essential gene prediction.

## I. INTRODUCTION

There are more than 20,000 genes of human being, which forms a redundant and high fault-tolerant system. Among these genes, some are vital for the survival and reproduction of us, but others are not. These two groups of genes are named as essential genes and nonessential genes. Essential genes are a group of fundamental genes necessary for a specific organism to survive in a specific environment. For human, essential genes refer to a subset of genes that are indispensable for the viability of individual human cell types [1][2]. These essential genes encode conservative functional elements which mainly contribute to DNA replication, gene translation, gene transcription, and substance transportation. The identification and analysis of essential genes is very important for understanding the minimal requirements of basic life, and it’s vital for drug-target identification, synthetic biology, and cancer research.

There are two ways to identify essential genes, wet lab experimental methods and computational methods. For example, gene direct deletion and transposon-based randomized mutagenesis have been used to identify essential genes for bacteria and yeast in the genome scale [3]; microinject KO and nuclear transfer techniques have been used to identify essential genes in mice [4]; the CRISPR/Cas9 genome editing system has been used to identify essential genes from human cell lines [1][2][5]. Experimental methods are often costly, time-consuming, and laborious. The accumulation of essential gene datasets and the sequence data as well as multiple functional data enables researchers to explore the relationships between gene essentiality and different genomics data, and to develop effective models to predict gene essentiality. These computational methods can greatly reduce the cost and time involved in for finding essential genes which further boosts our understanding to basic life and to human diseases, and helps quickly finding new drug targets and developing drugs.

Computational methods can be classified into two groups. One focus is to design a centrality measure to rank proteins/genes, while the other focus is to integrate multiple features using machine learning to predict gene essentiality. The most widely used centrality measures include degree centrality, betweenness centrality, closeness centrality, eigenvector centrality, to name a few. These centrality measures have been found having relationship with gene essentiality in multiple model organisms and human [6][7], however they can only differentiate a subset of essential genes from nonessential genes. One reason might be the incompleteness and false positive/false negative interactions in the protein-protein interaction (PPI) networks, the other reason might be the fact that gene essentiality relates to multiple biological factors rather than only the topological characteristics. Due to these reasons, researchers have proposed some more complicated centrality measures by integrating network topological properties with other biological information. For example, Zhang et al. proposed a method, CoEWC, to capture the common properties of essential genes in both date hubs and party hubs by integrating network topological property with gene expression profiles, which showed significant improvement of prediction ability compared to those only based on PPI networks [6]. An ensemble framework was also proposed based on gene expression data and PPI network, which can greatly enhance the prediction power of common used centrality measures [8]. Luo et al. proposed a method LIDC to predict essential proteins by combining local interaction density with protein complexes [9]. Zhang et al. proposed OGN by integrating network topological properties, gene expression profiles, and orthologous information [10]. GOS was proposed by Li et al. by integrating gene expression, orthologous information, subcellular localization and PPI networks [11]. These proposed integrated centrality measures have improved prediction power over the ones only based on PPI network topological properties. However, they still have limited prediction accuracy since gene essentiality relates to many complicated factors which make it impossible to represent them by a scalar score. At this point, machine learning is a good choice to fully utilize multiple features for predicting essential genes.

Many machine learning models and deep learning frameworks have been proposed and successfully applied in different fields. For example, convolutional neural network (CNN) was used for biomedical image segmentation [12]; density-based neural network was used for pavement distress image classification [13]; CNN was also used for drug-target prediction [14]; active learning and transductive k-nearest neighbors were used for text categorization [15][16]; Support Vector Machines (SVM) was used for essential gene prediction [17][18]. In the research field of essential gene prediction, many machine learning based prediction methods have been proposed to integrate features from multiple omics data [19]. As shown in the review article [19], traditional machine learning methods were used to predict gene essentiality and most of them were evaluated on data from model organisms. In these methods, topological features together with features from sequence and other functional genomics data were extracted manually and then used to train the models. How to extract informative features is very important and challenging, which requires ample domain knowledge as well as the understanding of what relationship exist between gene essentiality and each omics data. Usually we only know that an omics data would contribute to gene essentiality, but we don’t know what attributes of it have such effect and how to represent it. This puts a limitation on traditional machine learning methods to obtain good prediction accuracy.

In recent years, deep learning techniques have been used to automatically extract features and to train a more powerful classification model for predicting essential genes. Grover et al. proposed a network embedding method based on deep learning, node2vec, to learn a low-dimension representation for each node [20]. This method has been used to extract topological features from PPI networks for predicting essential genes, and these features are more informative than those obtained via some popular centrality measures [7][21, 22]. CNN was used to extract local patterns from time-serials gene expression profiles from S. cerevisiae [21] and Zeng et al. also used bidirectional long short term memory (LSTM) cells to extract features from the same gene expression profiles [22]. The automatically learned features are combined with other manually extracted ones to train a deep learning model for human essential gene prediction [7]. A six hidden-layers neural network was designed to predict essential genes in microbes by only using manually extracted features from sequence data [23].

Recently, human essential genes have been identified in several human cancer cell lines by utilizing CRISPR-Cas9 and gene-trap technology [1][2][24]. These identified essential genes provide a clear definition of the requirements for sustaining the basic cell activities of individual human tumor cell types, and can be regarded as targets for cancer treatment. These essential gene datasets together with other available biological data sources enable us to test one important and interesting assumption that human gene essentiality might be accurately predicted using computational methods. In this paper, we propose a deep learning framework, DeepSF, to predict human essential genes. DeepSF integrates features derived from nucleotide sequence and protein sequence data, features learned from PPI network, features encoded using gene ontology (GO) enrichment scores, features from protein complexes, and features from protein domain data to train a multi-layers deep learning model. We show that DeepSF can accurately predict human essential genes.

## II. METHODS

DeepSF consists of two modules, feature extraction module and classification module. In its feature extraction module, it extracts/learns features from multiple omics data and integrates these features together as input to the classification module. In this paper, we mainly consider five types of features, while other types of features can be easily integrated into our model.

Features encoded with gene ontology (GO) enrichment scores are calculated as follows. We choose 100 GO terms from cellular component part to encode the genes. For each gene, we first obtain its direct neighbors from PPI network to form a gene set consisting of this gene and its neighbors, then do gene ontology enrichment analysis for this gene set against the 100 GO terms using hypergeometric test. The enrichment score is calculated as −log10(p-value) for each GO term. In this way, we get a 100-dimension feature vector for each gene.

From protein complexes, we extract two features for each gene. The first feature is the number of protein complexes a gene involved in. The second feature is calculated as follows. For a gene, we first get the gene set N consisting of its direct neighbors in a PPI network. Suppose there are M neighbors of this gene involved in a protein complex. We calculate a score s = |M|/|N| as the ratio of its neighbors involved in a protein complex. The second feature is the sum of the ratios across all the protein complexes.

Network features are learned based on a network embedding method, node2vec. Each gene is represented by a 64-dimension feature vector learned from the PPI network. Previous studies showed that this low-dimension representation learned using node2vec is superior than the features calculated by popular centrality measures [7, 21, 22].

Sequence features consist of codon frequency, maximum relative synonymous codon usage (RSCUmax), codon adaptation index (CAI), gene length, GC content, amino acid frequency, and protein sequence length. There are 89 sequence features in total. For more details about how these features are calculated, please refer to [7].

From protein domain data, we extract three features for each gene. The first feature is the number of domain types a protein has. The second feature is the number of unique domain types a protein has. The third feature is the sum of inverse domain frequency (IDF). The frequency f of a domain is the number of proteins which have this domain. It’s IDF is 1/f. Suppose a protein u has n domains, then the third feature of u is calculated as in (1).

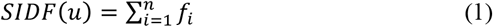

The classification module of DeepSF is based on the multilayer perceptron structure. It includes one input layer, three hidden layers, and one output layer. We use rectified linear unit (ReLU) as the activation function for all the hidden layers, while the output layer uses sigmoid activation function to perform discrete classification. The loss function in DeepSF is binary cross-entropy. A dropout layer is followed each hidden layer to make the network less sensitive to noise in the training data and to increase its generalization ability. In order to address the imbalanced learning issue inherent in essential gene prediction problem, we utilize class weight to train a weighted neural network, which will allow the model to pay more attention to instances from minority class than those from majority class and give larger penalty when it misclassify an instance from minority class. This way urges it to learn a more balanced and effective classifier.

## III. RESULTS AND DISCUSSION

### A. DATASETS

Human essential genes are downloaded from DEG database [25]. There are 16 human essential gene datasets among which 13 datasets are from [1][2][24]. We choose the genes contained at least in 5 datasets as our essential gene dataset. Excluding the genes annotated to essential genes in DEG, the other genes are considered as nonessential genes.

The DNA sequence data and protein sequence data are downloaded from Ensembl [26] (release 97, July 2019). We download the PPI data from BioGRID [27] (release 3.5.181, February 2020). Only physical interactions between human genes are used. After filtering out self-interactions and several small separated subgraphs, we obtain a protein-protein interaction network with 17,762 nodes and 355,647 edges. This interaction network is used to learn embedding features for each gene. It’s also used in aiding to compute some features from gene ontology, protein complex, and protein domain.

Gene Ontology data are downloaded from Gene Ontology website [28][29] and protein complex data are downloaded from CORUM [30]. Protein domain data are from Pfam database [31], and we collect this data via ensembl website [26]. The genes having sequence features, network embedding features as well as GO enrichment scores are used for the following classification performance evaluation. In total 2009 essential genes and 8414 nonessential genes are used for the following analysis.

### B. EVALUATION METRICS

We use multiple metrics to evaluate the performance of DeepSF. The first metric is the area under the receiver operating characteristic (ROC) curve (AUC). ROC plot represents the trade-off between sensitivity and specificity for all possible thresholds. The second metric is the area under the precision-recall curve (auPRC). Precision-Recall (PR) curves summarize the trade-off between the true positive rate and the positive predictive value using different probability thresholds. ROC curves are appropriate for balanced classification problems while PR curves are more appropriate for imbalanced datasets. Since essential gene prediction here is an imbalanced classification problem, the auPRC metric is more important than AUC. The third metric is Matthews correlation coefficient (MCC) [32], and the fourth metric is F1 measure. In addition to these four comprehensive metrics, we also give the following performance measures: sensitivity (Sn), specificity (Sp), positive predictive value (PPV), and accuracy (Ac), which are defined in (2) - (5), MCC and F1 in (6) and (7), where TP, TN, FP, and FN are the number of true positives, true negatives, false positives, and false negatives, respectively.

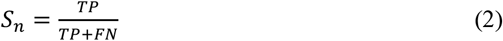

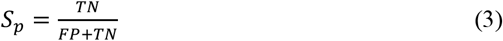

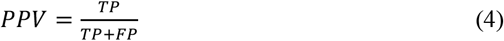

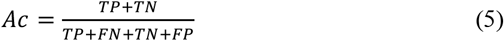

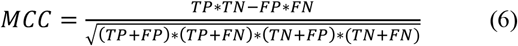

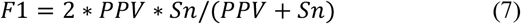

### C. PERFORMANCE EVALUATION

In the following experiments, we use the parameters for DeepSF as follows. Adam with initial learning rate 0.001 is used as the optimizer. The sizes for hidden layers are 1024, 512, 256. The dropout rate is 0.2. The training runs for 200 epochs using early stopping criteria with patience = 15. In order to cope with the imbalanced learning issue, we set class weight to 4.5 for essential genes, and 1 for nonessential genes. The stratified randomized 10-fold cross validation is used to evaluate the performance of DeepSF. At each fold, 10% data are held out for testing, and 90% data are used for training among which 10% is used as the validation data in the training process.

Figure 1 presents the ROC and PR curves of DeepSF across the 10-fold cross validation. From figure 1 we can see that DeepSF reaches its best performance at fold 7 with AUC = 0.97 and auPRC = 0.94. The average AUC of DeepSF across the 10-fold cross validation is 95.17% with standard deviation STD = 0.65%, and the average auPRC is 92.21% with STD = 1.37%. In addition, the performance of DeepSF is quite stable since the difference is about 2.3% (4%) between its best and worst AUC (auPRC) scores across the 10-fold cross validation. The worst auPRC score is still above 90% which indicates that DeepSF is very effective for predicting human essential genes. In addition to good scores of AUC and auPRC, its average accuracy, MCC and F1 scores are 91.59%, 73.57%, and 78.71% respectively.

**FIGURE 1.**
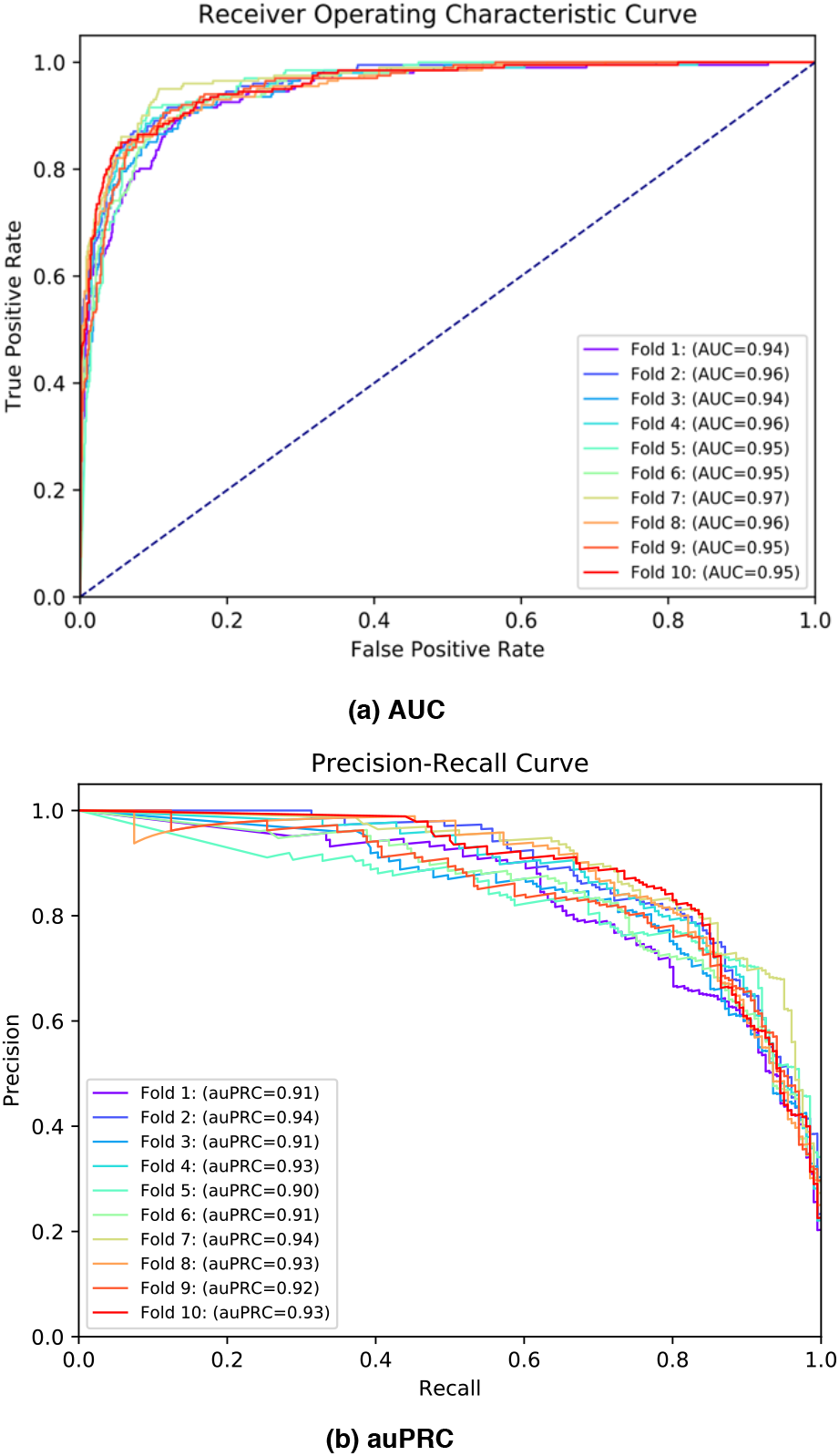
The ROC and PR curves of DeepSF.

### D. PERFORMANCE COMPARISON WITH OTHER MACHINE LEARNING MODELS

In order to demonstrate the superior of DeepSF, we also compare it with three popular traditional machine learning models (SVM, RF, and Adaboost) as well as a recent deep learning based model DeepHE. For fair comparison, DeepSF and the three traditional machine learning models use the same input features so that the only difference here is the classification module. DeepSF uses deep learning techniques while the three methods are all traditional machine learning models. For DeepHE, we use the features presented in [7] and set the hidden layers to 128, 256, 512 as used in [7] while keeping the training data and other settings same with DeepSF.

Figure 2 shows the box plot for the performance comparison between DeepSF and the four compared models across 10-fold cross validation. From figure 2 we can see that the AUC and auPRC scores of DeepSF are significantly higher than those of the three traditional machine learning models, and are slightly better than those of DeepHE. The average AUC of DeepSF increases 1.73%, 7.3%, 22.81%, and 13.58% compared with that of DeepHE, SVM, RF, and Adaboost respectively. The average auPRC of DeepSF increases 2.99%, 41.12%, 61.73%, and 48.23% compared with that of DeepHE, SVM, RF, and Adaboost respectively. In terms of the other two comprehensive metrics MCC and F1, DeepSF achieves similar performance with SVM, but obvious better than DeepHE, RF and Adaboost. For example, the average MCC of DeepSF increases 7.24%, 0.07%, 14.27%, and 5.39% compared with that of DeepHE, SVM, RF, and Adaboost respectively. The average F1 of DeepSF increases 5.41%, - 0.04%, 14.98%, and 4.55% compared with that of DeepHE, SVM, RF, and Adaboost respectively. From the above experimental results and the fact that auPRC is the most suitable metric for imbalanced classification problem, we can conclude that our deep learning based model is superior than the traditional machine learning models and that integrating suitable features from more biological data sources can improve the prediction performance.

**FIGURE 2.**
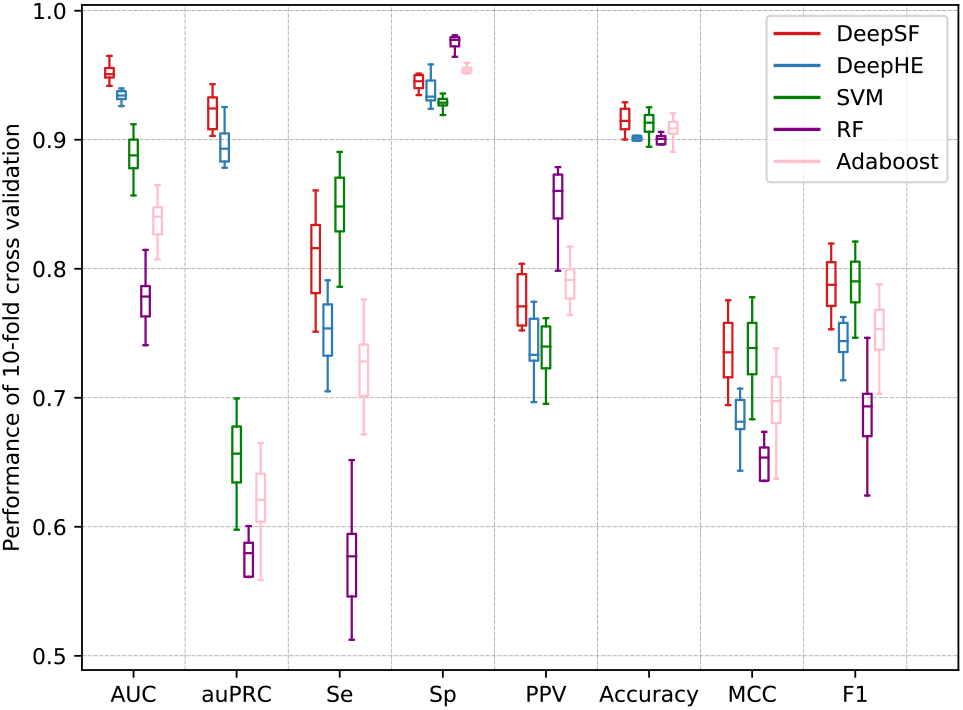
Performance comparison of DeepSF, DeepHE, SVM, RF, and Adaboost.

### E. ABLATION STUDY

In order to see the contribution of each type of features used in DeepSF, we also evaluate it by removing one type of features each time. The experiments use same settings except the input features. Figure 3 gives the box plot for DeepSF with different types of input features, which tells us that DeepSF with the integration of all the five types of features performs best which further confirms the contribution and complement of these five types of features. By removing one type of features each time, we can evaluate the contribution of each type of features to the overall performance. From figure 3 we can see that the combination of N + G + C + D performs worst which might be due to the fact that among which three types of features utilize the PPI network topological information so that they have less complement effect. The other four combinations with sequence features perform only slightly worse than that of using all the five types of features, which tells us that sequence features have big complement with the other types of features. As shown in [7], when only using one type of features, deep model using network embedding features performs better than it using sequence features. Therefore, the above phenomenon doesn’t indicate that sequence features are superior than network features, but that sequence features are more complementary to other three types of features.

**FIGURE 3.**
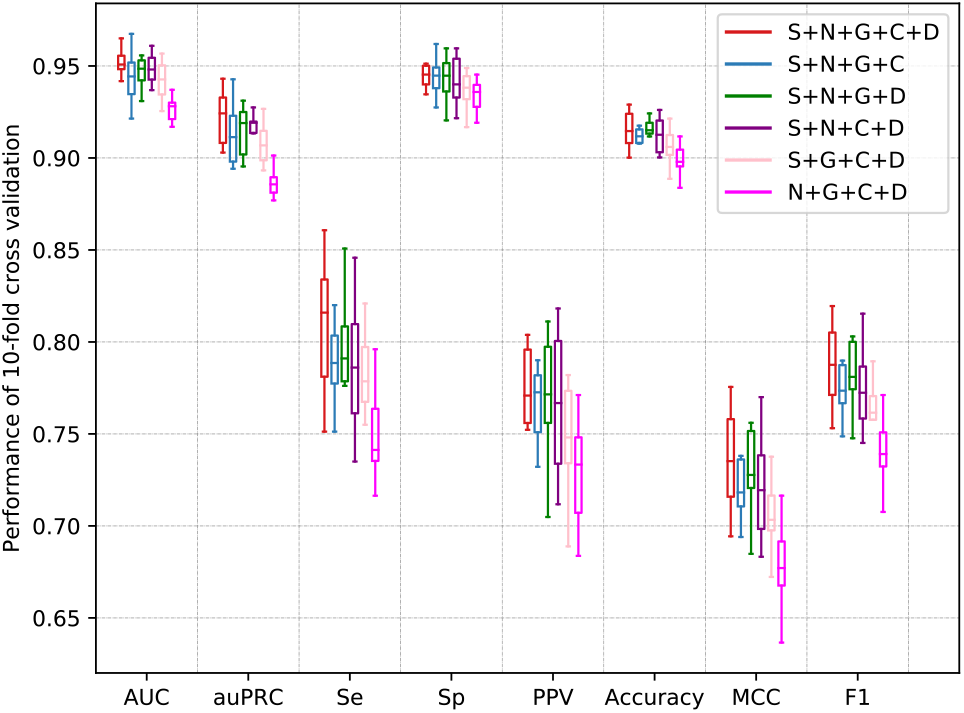
Performance comparison of DeepSF with different types of features. S: sequence features; N: network embedding features; G: gene ontology features; C: protein complex features; D: protein domain features.

## IV. CONCLUSIONS

In this paper, we propose a deep learning based method, DeepSF, to predict human essential genes. DeepSF integrates five types of features extracted/learned from sequence and functional genomics data, and utilizes deep learning techniques to train a cost-sensitive classifier to enhance prediction accuracy of gene essentiality. Our 10-fold cross validation experiments demonstrate that deep learning is more superior than the traditional machine learning models, and that the integration of features from more different omics data can enhance the prediction accuracy. Our experiments demonstrate that DeepSF is effective for predicting human essential genes.

In the future, we are interested in how to use deep learning to automatically learn features from different types of biological data. For example, how to learn a low-dimension representation if we use all GO terms to encode genes instead of the subset of selected GO terms. Exploring and integrating more biological data into the learning and classification model is another interesting direction, such as epigenomics data and gene expression profiles. Predicting cancer cell line specific essential genes by designing effective deep learning models and integrate cell line specific information would be very interesting. In addition, we are also interested in testing whether data editing [33] and clustering aided techniques [34] are useful for imbalanced learning problem. Membrane computing is a distributed parallel computing paradigm [35] and membrane computing models have great potential in solving fault diagnosis [36] as a kind of machine learning models and learning problem of models [37]. We are also interested in exploring the benefit of membrane computing models to improve the prediction performance of gene essentiality.

---

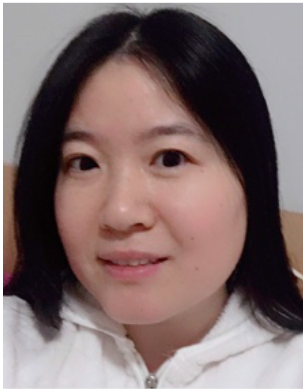

**Xue Zhang** received the PhD degree in Computer Science from Southeast University, China, in 2007. Then she joined Peking University as a postdoc fellow focusing on data mining, machine learning, and bioinformatics. From 2014 to 2016, she worked at National Institutes of Health (NIH/NHLBI) as a research scientist. She is currently working at Tufts University School of Medicine as a research associate. Her current research interests include network biology, bioinformatics, machine learning and deep learning.

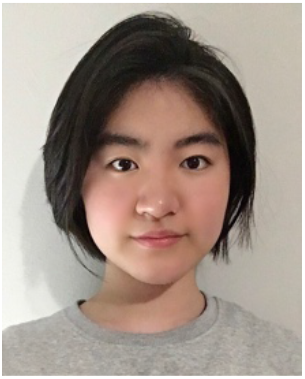

**Weijia Xiao** is currently a senior at Boston Latin School, Boston, USA. She got secondary award in MIT BWSIX RACECAR final competition in 2020. She also ranked 2^nd^ place in AIHacks II Hackathon in 2020. Her interests include machine learning, deep learning, artificial intelligence, and bioinformatics.

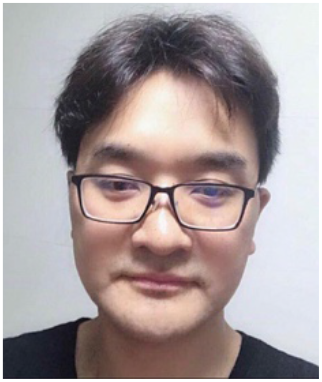

**Wangxin Xiao** received the PhD degree in Traffic Information Engineering from Southeast University, China, in 2005. Then he joined Wuhan University of Technology as a postdoc fellow focusing on data mining and pattern recognition from 2005 to 2008. He is currently a professor at Huiyin Institute of Technology. His research interests include image processing, machine learning and bioinformatics.

